# Integrative analysis of transcriptomic and clinical data uncovers the tumor suppressive activity of MITF in prostate cancer

**DOI:** 10.1101/391763

**Authors:** Lorea Valcarcel-Jimenez, Alice Macchia, Natalia Martín-Martín, Ana Rosa Cortazar, Ariane Schaub-Clerigué, Mikel Pujana-Vaquerizo, Sonia Fernández-Ruiz, Isabel Lacasa-Viscasillas, Aida Santos-Martin, Ana Loizaga-Iriarte, Miguel Unda-Urzaiz, Ivana Hermanova, lanire Astobiza, Mariona Graupera, Julia Starkova, James Sutherland, Rosa Barrio, Ana M. Aransay, Arkaitz Carracedo, Verónica Torrano

## Abstract

The dysregulation of gene expression is an enabling hallmark of cancer. Computational analysis of transcriptomics data from human cancer specimens, complemented with exhaustive clinical annotation, provides an opportunity to identify core regulators of the tumorigenic process. Here we exploit well-annotated clinical datasets of prostate cancer for the discovery of transcriptional regulators relevant to prostate cancer. Following this rationale, we identify Microphthalmia-associated transcription factor (MITF) as a prostate tumor suppressor among a subset of transcription factors. Importantly, we further interrogate transcriptomics and clinical data to refine MITF perturbation-based empirical assays and unveil Crystallin Alpha B (CRYAB) as an unprecedented direct target of the transcription factor that is, at least in part, responsible for its tumor suppressive activity in prostate cancer. This evidence was supported by the enhanced prognostic potential of a signature based on the concomitant alteration of MITF and CRYAB in prostate cancer patients. In sum, our study provides proof-of-concept evidence of the potential of the bioinformatics screen of publicly available cancer patient databases as discovery platforms, and demonstrates that the MITF-CRYAB axis controls prostate cancer biology.

## Introduction

Balanced integration of intracellular circuits operates within a normal cell to sustain physiological homeostasis. Alterations in some, if not all, of these circuits converge in changes on gene expression, which will eventually enable the acquisition and sustenance of the hallmarks of cancer cells (1). This event emphasizes the importance of maintaining the transcriptional homeostasis in normal cells and places gene expression deregulation at the core of cancer research interests.

In the last decades, transcriptomics data derived from cancer specimens has become an important resource for the classification, stratification and molecular driver identification in tumors. We and others have demonstrated that deregulation of gene expression is a key node for cancer pathogenesis and progression (2–6). Prostate cancer (PCa) research exemplifies the effort in deciphering the genomics and transcriptomics landscape of tumors, and extremely valuable data has been generated (7–13). In spite of the public availability of these relevant data, they are still underexploited by the scientific community to understand PCa biology. In this regard, the computational tools and dataset selection strategies to carry out these studies are a bottleneck for the cancer research field.

By combining integrated-bioinformatics screening of clinically relevant PCa datasets with *in vivo* and *in vitro* molecular biology assays, we have recently described the metastasis suppressor activity of Peroxisome proliferator-activated receptor *γ* (PPAR*γ*) coactivator alpha (PGC1α) (14, 15). This transcriptional coactivator is a major regulator of mitochondrial biogenesis and function, and has an inherent capacity to integrate environmental signals and cellular energetic demands. This ability empowers PGC1 α to be a driver in shaping responses to metabolic stress during different physiologic and tumorigenic processes (16). As might be expected due to its fundamental role in normal and cancer scenarios, the regulation of PGC1α expression, from the genomic to the protein level, is complex and dynamic (17). At the level of mRNA expression, one of the well-defined direct regulators of PGC1ais the Microphthalmia-associated transcription factor (MITF) (18).

MITF is a basic helix-loop-helix leucine zipper (bHLHZIP) transcription factor that regulates the expression of lineage commitment programs that are essential for propagation of the melanocyte lineage (19). The existence of different MITF transcript variants is the result of both alternative splicing and promoter activation that results in the cell-type-specific expression of the different MITF isoforms (A, CX, MC, C, E, H, D, B, M, J) (20). The melanoma specific isoform M-MITF is the best studied isoform and, despite some controversy, its expression is generally deregulated in melanoma. Although MITF alone cannot act as a classical oncogene, it has been called a ‘lineage survival oncogene’ for melanoma (19, 21). Importantly, the presence or absence of the M-MITF-PGC1α regulatory axis has stratification potential in melanoma and informs on the efficacy of BRAF inhibitor treatments (18, 22). Although the expression of MITF has been detected in other types of tumors different from melanoma (23, 24), its active role in the progression of these diseases, including PCa, remains unexplored.

Crystallin Alpha-B (CRYAB) is a ubiquitous small heat shock protein that is expressed in response to a wide range of physiological and non-physiological conditions preventing aggregations of denatured proteins. In a wide variety of tumor types CRYAB has been found to be overexpressed and associated with disease progression (25–29) and poor prognosis (30, 31). However, in PCa and nasopharyngeal cancers CRYAB expression is decreased (32, 33), pointing at possible tumor suppressive activity of CRYAB in these cancer scenarios.

In the present study, by combining an exhaustive interrogation of seven publically available prostate cancer databases with refined empirical assays, we have identified MITF as a prostate tumor suppressor. In addition, we have unveiled CRYAB as a novel direct target of the transcription factor that is, at least in part, responsible for its tumor suppressive activity in prostate cancer. Importantly, the tumor suppressive role for this novel MITF-CRYAB axis is supported by the enhanced prognostic potential of a signature based on the concomitant alteration of both genes in PCa patients.

## Results

### Bioinformatics screening identifies MITF as a transcription factor altered in prostate cancer

We have recently demonstrated that the reduced expression of the transcriptional co-activator PGC1α is a causal event for metastatic prostate cancer (PCa) (14). We sought to identify transcriptional regulators related to the alteration in PGC1 α expression. We designed a bioinformatics strategy based on the analysis of 16 genes directly linked to the regulation of *PGC1A* gene (17, 22, 34–38), in order to identify transcription factors that could be relevant to PCa biology. For the candidate screen we applied selection criteria based on the consistency of, first, the correlation with *PGC1A* expression and second, the expression of each individual candidate in seven publicly available PCa datasets (7, 9–13) (Figure 1A). We selected those candidates whose expression in primary tumors correlated with *PGC1A* (R ≥ 0.2 and p-value ≤ 0.05 in more than 50% of the datasets) (Supplementary Figure 1 A) and was altered when compared to normal specimens. For genes exhibiting various transcript variants, the correlation analysis was initially performed using the average signal (Supplementary Figure 1 A) and, when available (only Taylor dataset (11)), the correlation was confirmed in all the individual isoforms (Supplementary table 1). The transcription factor MITF was the sole candidate that complied with the established criteria. We observed a consistent correlation between *PGC1A* and *MITF* in four out of the seven datasets analyzed (Figure 1 B and Supplementary Figure 1 A). In addition, not only the mean expression but also the expression of the individual *MITF* isoforms were reduced in primary and metastatic PCa specimens when compared with the normal prostate tissue samples (Figure 1 C and Supplementary Figure 1 B). Taken together, our data reveals MITF as a PGC1A-associated transcription factor that is consistently downregulated in PCa.

**Figure 1.**
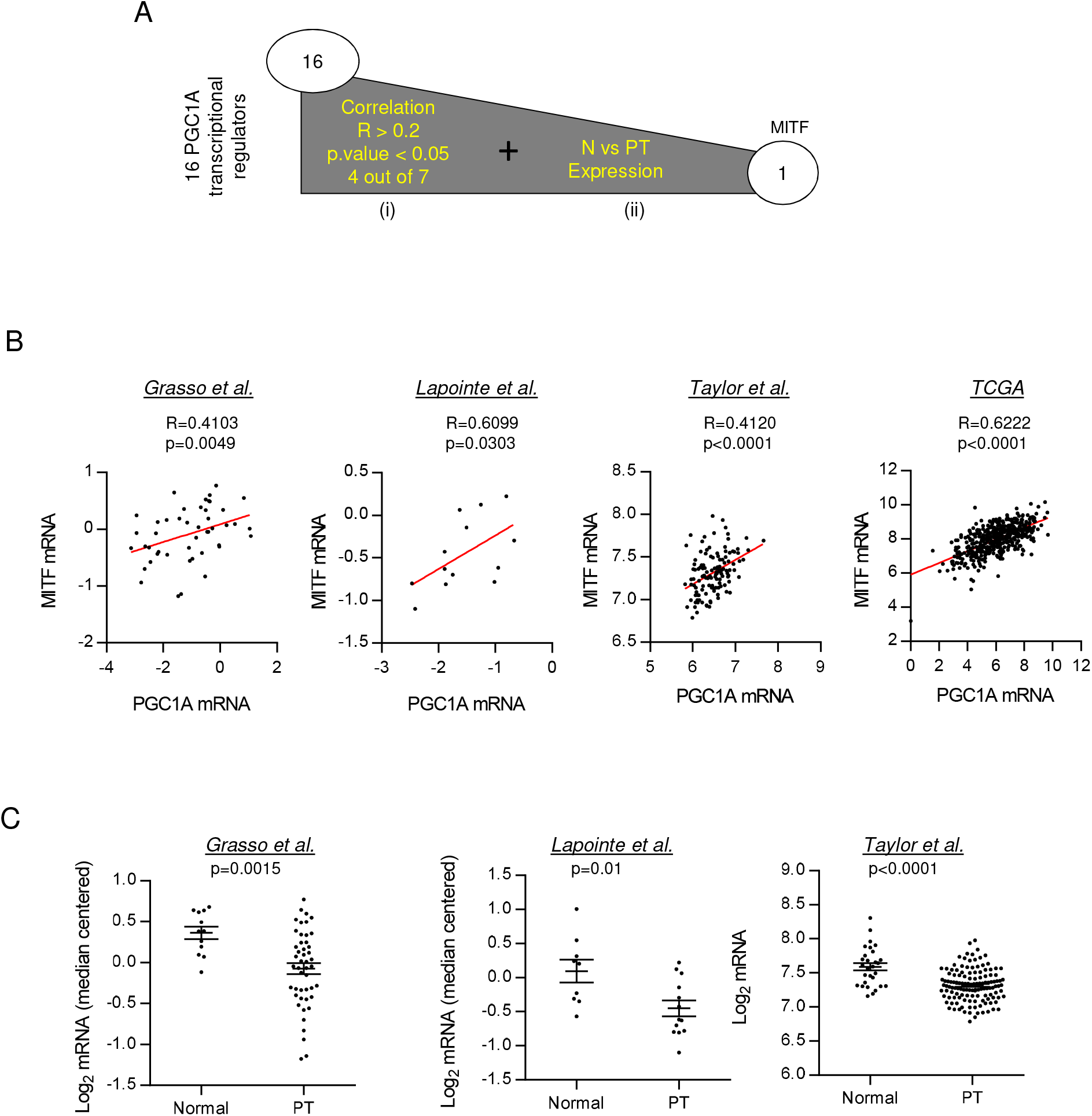
MITF expression correlates with PGC1A expression and is downregulated in PCa. **A.** Schematic representation of candidate screening to mediate PGC1A downregulation in PCa. Candidate selection was performed by applying two different selection criteria based on the consistency within the datasets used (>50%): the expression of the candidate must be consistently (i) correlated with the PGC1A’s and (ii) altered in the disease. **B.** Correlation analysis between PGC1A and MITF expression in primary tumor (PT) specimens of different PCa datasets ((9–11) and TCGA provisional). Sample sizes: Grasso n=45; Lapointe n=13; Taylor n=131 and TCGA provisional n=495. **C.** MITF expression in normal prostate and primary tumors (PT) specimens in different data sets (9–11). Correlation (B) and expression (C) data from Taylor dataset corresponds to the mean signal of all isoforms of the transcripts. In B and C, each dot corresponds to an individual specimen. Sample sizes: Grasso et al. (Normal, n=12; PT, n=45); Lapointe et al. (Normal, n=9; PT, n=13); Taylor et al. (Normal, n=29; PT, n=131). Error bars represent s.e.m. Statistic test: Spearman correlation *R* (B) and Mann Whitney test (C). p, p-value.

### MITF exhibits tumor suppressive activity in PCa

The expression profile of *MITF* in PCa, together with its direct correlation with *PGC1A,* was suggestive of a tumor suppressive activity of the transcription factor. We first examined the differential expression of the distinct mRNA isoforms of *MITF* in normal, PCa primary tumors and PCa cell lines (Supplementary Fig. 2 A-C). *MITFA* was the isoform predominantly expressed in the three scenarios analyzed, and we pursued the studies further with this isoform. Next, we aimed to analyze the biological consequences of ectopic expression of MITFA in PC3 PCa cells. We transduced PC3 cells with a lentiviral vector containing a doxycycline-inducible cassette for the expression of MITFA resulting in the generation of the PC3 TRIPZ-MITFA cell line. The induction of MITFA expression (Figure 2 A and B) as well as the regulation of known target genes, including *PGC1A* (14, 15) (Supplementary Figure 2 D-E) was confirmed. We next evaluated the biological outcome of MITFA ectopic expression in PC3 cells and observed that its upregulation significantly reduced two-dimensional and anchorage-independent growth (Figure 2 C and D), with no effect of doxycycline treatment by itself (14). In line with its known function as an inhibitor of cell cycle progression (39), the increased expression of MITFA in PC3 cells resulted in a decrease in BrdU incorporation, a surrogate readout of proliferation (Figure 2 E).

**Figure 2.**
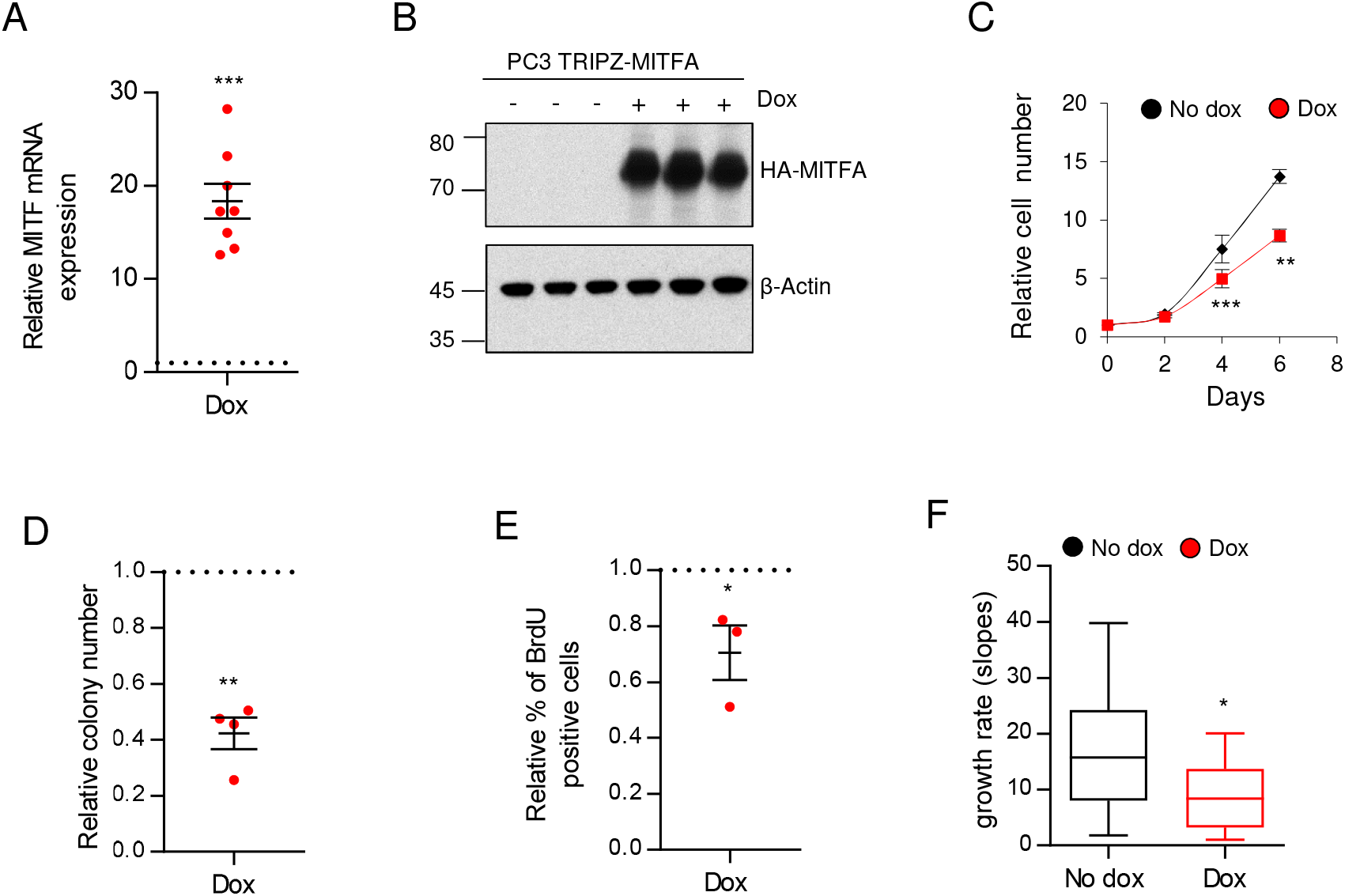
MITF exhibits tumor suppressive activity in PC3 PCa cell line. **A-B.** Analysis and quantification of MITF expression by qRTPCR (A, n=8)) and western blot (B, representative experiment out of 3 independent ones) in PC3 TRIPZ-MITFA cells after treatment with 0.5 μg mL^-1^ doxycycline (Dox). **C.** Relative cell number quantification by crystal violet in doxycycline treated and non-treated PC3 TRIPZ-MITFA cells. Data is normalized to day 0. Asterisks indicate statistics of 5 independent experiments. One representative experiment out of 5 is shown. Error bars represent standard deviation. **D-E.** Effect of MITF induction on anchorage independent growth (D, soft agar; n=4 independent experiments) and BrdU incorporation (E, n=3 independent experiments). **F.** Impact of MITF induction in tumor growth rate of PC3 TRIPZ-MITFA cells (n=7 animals per group; 14 injections/tumors). No dox: MITFA non-induced conditions; Dox: MITFA induced conditions. Error bars represent s.e.m (A, D and E) or minimun and maximum values (H). Statistic test: One sample t-test (A, D and E) and Student t-test (C and F. *p < 0.05, **p < 0.01, ***p < 0.001.

In order to ascertain whether the regulation of endogenous *PGC1A* (Supplementary Figure 2 E) was required for the anti-proliferative effect of MITFA in PC3 cells, we aim at silencing *PGC1A* by using constitutive (pLKO) expression of short hairpins against it (Supplementary Figure 2 F). Transduction with the shRNA prevented the upregulation of *PGC1A* upon MITFA induction (Supplementary Figure 2 G) but the anti-proliferative effect of the transcription factor remained unaffected (Supplementary Figure 2 H). These data suggested that the reduced proliferation induced by MITFA was not dependent on the regulation of endogenous *PGC1A* in PC3 cells.

Importantly, the overall reduction in cell proliferation induced by MITFA was confirmed *in vivo.* Using subcutaneous xenografts assays we observed that MITFA over-expression in PC3 cells (Supplementary Fig. 2 I) led to a marked reduction in the tumor volume (Supplementary Fig. 2 J) and growth rate (Figure 2 F), with no changes in angiogenesis (Supplementary Figure 2 K). Altogether these results demonstrate that MITFA isoform exhibits tumor suppressive activity in PCa.

### Candidate screening of genes mediating the tumor suppressive activity of MITF

In order to decipher the molecular mechanism driving the tumor suppressive role of MITFA we performed gene expression profiling of both doxycycline treated and control PC3 TRIPZ-MITFA cells and identified 101 probes that showed statistically differential signal between both conditions (Supplementary table 2). We first performed a gene enrichment analysis with those genes which displayed upregulated expression (76 genes) upon MITFA over-expression (Figure 3 A and Supplementary table 3), as the number of downregulated genes (25) was no sufficient to obtain any gen enrichment. Next, we aimed at identifying potential MITFA effectors of relevance in human PCa. To this end, we established a threshold of 1.5 fold change over MITFA non-induced cells, which resulted in 8 probes (corresponding to 6 annotated genes) upregulated upon the induction of the transcription factor (Supplementary table 2; yellow bold highlighted). We next performed correlation analysis between *MITF* and each of the 6 differentially expressed genes obtained from the microarray (Figure 3 A and Supplementary Figure 3 A). The correlation analysis in PCa primary tumor specimens showed that a single gene, *Crystallin Alpha B (CRYAB),* had a consistent correlation (in more than 50% of datasets) with *MITF,* both the mean of isoforms (Figure 3 B and Supplementary Figure 3 A) and the individual isoform A (Supplementary table 4). The *MITF-CRYAB* correlation was confirmed using an independent cohort of PCa patients from a local hospital (Basurto cohort, Supplementary Figure 3 A). Moreover, the expression of CRYAB either at the level of mRNA (from public datasets and Basurto cohort) and protein (from Basurto cohort) was consistently downregulated through the progression of the disease (Figure 3 C and D and Supplementary Figure 3 B-D), supporting the association of MITF and CRYAB expression in PCa.

**Figure 3.**
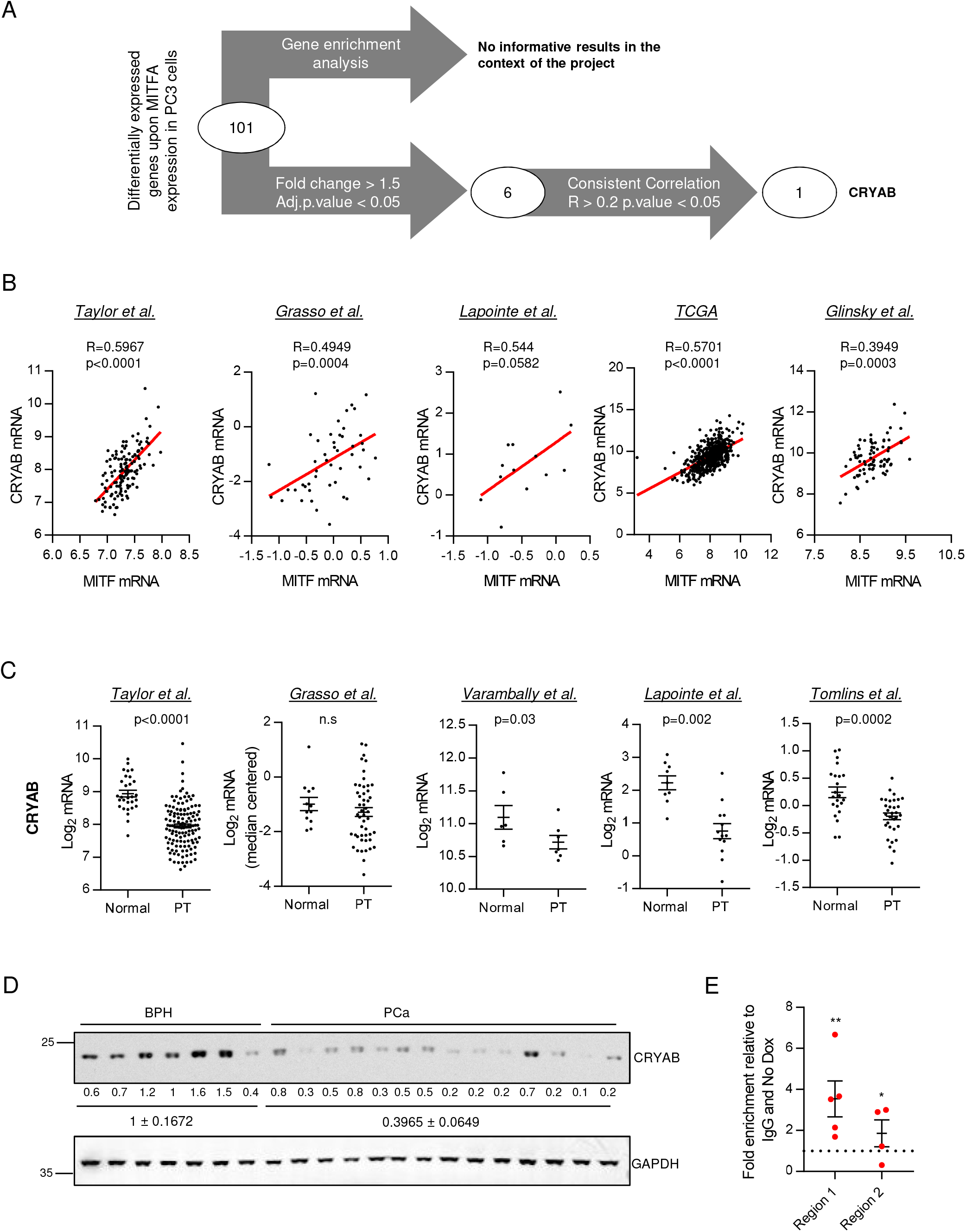
CRYAB is the candidate to mediate its tumor suppressive activity in PCa. **A.** Workflow of the candidate screening. **B.** Correlation analysis between MITF and CRYAB expression in primary tumor (PT) specimens of different PCa datasets. Sample sizes: Taylor, n=131; Grasso, n=49; Lapointe, n=13; TCGA provisional data, n=495; and Glinsky, n=78. **C.** CRYAB expression in normal prostate and primary tumor (PT) specimens in different PCa datasets (9–13). Sample sizes: Taylor (N, n=29; PT, n=130); Grasso (N, n=12; PT, n=49); Varambally (N, n=6; PT, n=7); Lapointe et al. (N, n=9; PT, n=13) and Tomlins (N, n=22; PT, n=32). Data from Taylor dataset corresponds to the mean signal of all isoforms of the transcripts. In B and C, each dot corresponds to an individual specimen. D. Western blot analysis of CRYAB expression in benign prostatic hyperplasia (BPH) and PCa specimens from Basurto University Hospital cohort (BPH n=7 patient specimens; PCa n=14 patient specimens). **E.** Chromatin immunoprecipitation (ChIP) of exogenous MITF on CRYAB promoter in PC3 TRIPZ-MITFA cells after induction with 0.5 μg mL-1 doxycycline for 3 days (n=4-5). Binding to ANGPT4 was used as a negative control. Final data was normalized to IgG (negative-immunoprecipitation control) and to No dox condition. No dox: MITFA non-induced conditions; Dox: MITFA induced conditions. Statistic tests: Spearman correlation (B); Mann Whitney test (C); one sample t test (E); Error bars represent s.e.m. *p < 0.05, **p < 0.01.

The regulation of CRYAB expression by MITFA was further validated *in vitro* by western blot and quantitative real-time PCR (qRTPCR) in doxycycline-treated PC3 TRIPZ-MITFA cell lines and *in vivo* by qRTPCR in the xenograft samples (Supplementary Figure 3 E-G). MITF is a transcription factor that regulates gene expression through the DNA binding to E-boxes (Myc-binding sites) (19). In order to confirm the direct regulation of *CRYAB* expression by MITFA, we screened the promoter of the chaperon and performed chromatin immunoprecipitation assays in two Myc-binding sites (UCSC-Genome browser; Supplementary Figure 3 H). As predicted, upon doxycycline treatment we detected differential binding of MITFA in both regions of *CRYAB* promoter (Figure 3 E).

Taken together, these data presented *CRYAB* as a direct target of MITFA and the best candidate to mediate its tumor suppressive activity in PCa.

### CRYAB mediates the tumor suppressive activity of MITF in PCa

We next studied the functional relevance of CRYAB for the tumor suppressive activity of MITFA in PCa. Towards this aim, we constitutively silenced the expression of *CRYAB* by RNAi using two independent short hairpin RNA (sh#1 and sh#2) in PC3 TRIPZ-MITFA cells. After validation that RNAi was achieved (Figure 4 A and Supplementary Figure 4 A-C) the tumor suppressive activity of MITFA was monitored in control and CRYAB-silenced conditions (PC3 TRIPZ-MITFA scr, sh#1 or sh#2 cell lines). *CRYAB* silencing blunted the anti-proliferative effects of MITFA *in vitro* in two-dimensional and anchorage independent growth when compared with scramble shRNA (Figure 4 B and C). Moreover, the reduction in BrdU induced by MITFA was prevented when *CRYAB* was silenced (Figure 4 D). Importantly, the requirement of CRYAB for the tumor suppressive activity of MITFA was corroborated *in vivo* (Figure 4 E and Supplementary Figure 4 D-F). The *in vitro* and *in vivo* data demonstrate that the induction of CRYAB is a major effector involved in the tumor suppressive activity of the transcription factor MITF in PCa.

**Figure 4.**
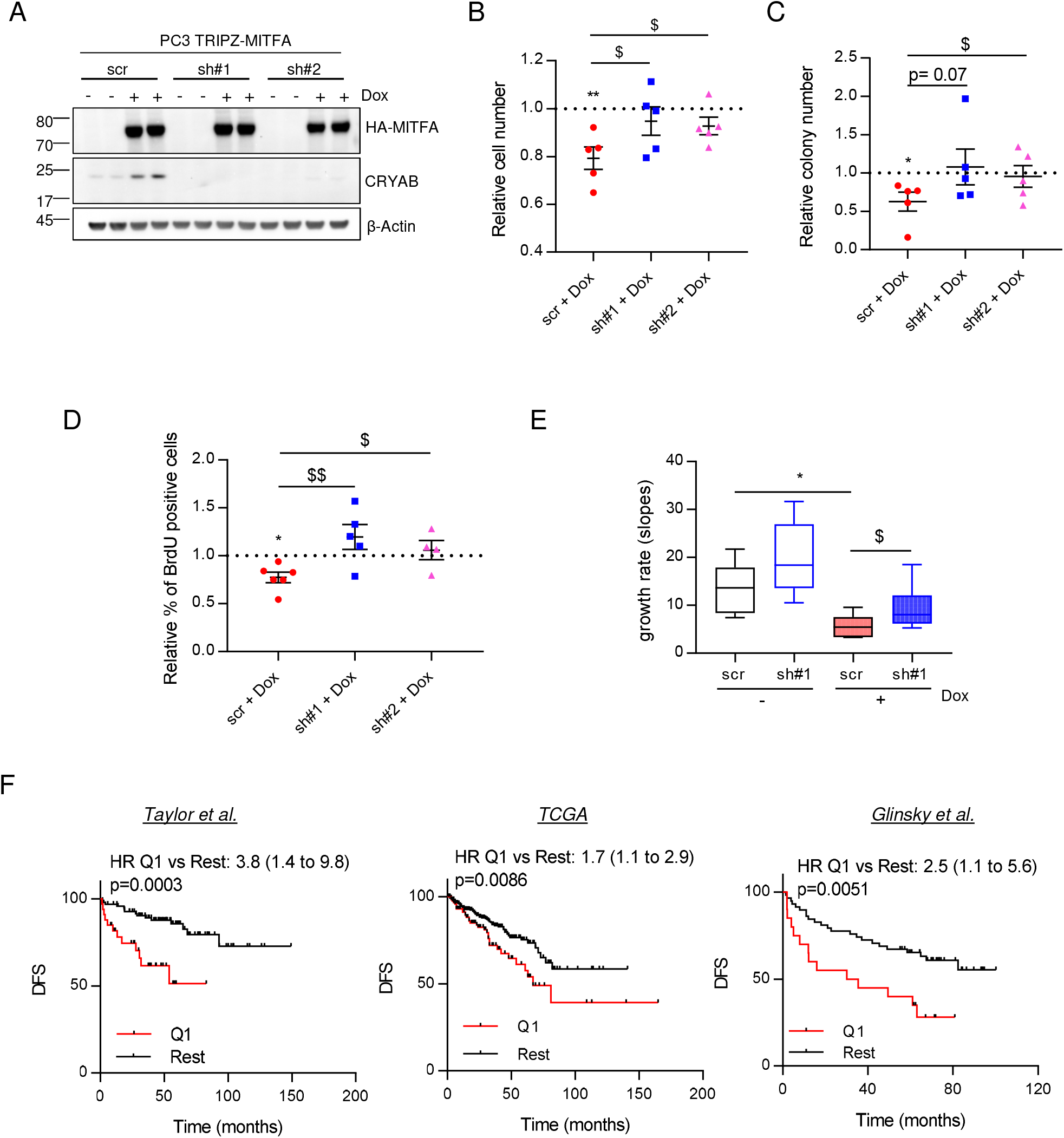
CRYAB mediates the tumor suppressor activity of MITF. **A.** Analysis of MITFA and CRYAB protein expression in doxycycline-treated PC3 TRIPZ-MITFA cells transduced with shScramble (scr) or two independent shCRYAB (sh#1 and#2) (one representative experiment with technical duplicates is shown; similar results were obtained in three independent experiments). **B.** Relative cell number quantification by crystal violet in doxycycline-treated and non-treated PC3 TRIPZ-MITFA cells, in the presence (scr) or absence (sh#1, 2) of CRYAB (n=5 independent experiments). Data is normalized to day 0 and represented as cell number at day 6 relative to No Dox condition (depicted by a dotted line). **C-D.** Effect of CRYAB silencing on anchorage independent growth (C, soft agar; n=4 independent experiments) and BrdU incorporation (D, n=3 independent experiments) in PC3 TRIPZ-MITFA cells after treatment with 0.5 μg mL^-1^ doxycycline. **E.** Impact of CRYAB silencing in tumor growth rate of MITF-induced cells (n=10 animals per group-scr or sh#1; 2 injections per mice; (scr No dox, n=10 tumors; sh#1 No dox, n=8 tumors; scr Dox, n=6 tumors; sh#1 Dox, n=11 tumors). **F.** Association of the mean signal of MITF and CRYAB with disease-free survival (DFS) in three PCa data sets (Q1: first quartile distribution; rest: second, third and fourth quartile distribution. Sample sizes: Taylor, primary tumors n=131; TCGA provisional data primary tumors n=490; Glinsky, primary tumors n=78. No dox: MITFA non-induced conditions; Dox: MITFA induced conditions. HR: Hazard Ratio. Statistic tests: One sample t test (B, C and D – No dox vs Dox conditions); Unpaired Student t-test (*t*) (B, C and D – Dox-treated scr vs Dox-treated sh#1/2); Log-rank (Mantel-Cox) test (F). Error bars represent s.e.m. */$ p < 0.05, **/$$p < 0.01. Asterisks indicate statistic between No dox and Dox conditions and dollar symbol between Dox-treated scr and Dox-treated sh#1 or 2.

We next asked whether the functional association between MITF and CRYAB could be employed to identify PCa patients with high disease aggressiveness. We thus ascertained the stratification potential of the MITF-CRYAB axis in PCa by means of consistency and robustness. We download the mRNA expression raw data together with the clinical data (recurrence or not recurrence) from Taylor (11), Glinsky (8) and TCGA (7) datasets. The individual or average expression signal of *CRYAB* and *MITF* genes was calculated for each patient in each dataset. Patients were separated by quartiles according to the individual or average signal of *CRYAB* and *MITF* genes and then Kaplan-Meyer survival curves were plotted comparing patients with low expression (Quartile 1 – (Q1)) of the individual genes or the gene combination (*CRYAB* and *MITF) versus* the rest of the cohort (Q2+Q3+Q4). Strikingly, the signature formed by the average signal of *MITF* and *CRYAB* outperformed the prognostic potential of each individual gene, strongly suggesting that the pathway described herein is strongly associated to PCa aggressiveness (Figure 4 F and Supplementary Figure 5).

Our data provide solid evidence of an unprecedented MITFA-CRYAB transcriptional axis that exerts tumor suppressive activity in PCa and could positively contribute to disease prognosis.

## DISCUSSION

Technological advances in the molecular understanding of cancer have led to a paradigmatic change in the way that we combat the disease. We are now able to deconstruct a tumor at a molecular level using genomics, transcriptomics, proteomics and metabolomics. This, in turn, enables us to foresee, identify and demonstrate the potential of patient stratification. Specifically, the transcriptomics characterization of tumors is an invaluable strategy to identify clinically relevant genes that play key roles in the progression of cancer, especially for those types with poorer prognosis (14). Thus, the comprehensive and integrative analysis of gene expression changes and clinical parameters in cancer has become a mainstream in cancer research. Mining cancer-associated transcriptome datasets is an emerging approach used by top cancer research groups, but better tools are needed to increase its power and user-friendliness. In order to face this challenge, new interfaces to exploit OMICs data, such as cBioportal (40, 41) are being designed to help scientists interrogate, integrate and visualize large amount of information contained on multiple credible and qualified cancer datasets.

In the present study, we exploited publicly available and well-annotated (transcriptomics and clinical data) prostate cancer databases together with experimental assays to describe a novel tumor suppressive activity of the transcription factor MITF in PCa, which is executed, at least in part, through the direct regulation of the CRYAB expression.

The functional implication of MITF in cancer has been best defined in melanoma, in which the expression of the transcription factor is heterogeneous. Although some controversy exists regarding its oncogenic role in melanoma, MITF has been defined as a “lineage survival oncogene” with no data pointing out at a tumor suppressive function (19, 21, 39, 42–45). Even though the expression of MITF has been detected in other cancer types (23, 24, 46), no data supporting a functional role of MITF deregulation has been reported yet in a cancer scenario different from melanoma.

Here we show that MITF is downregulated in PCa when compared with normal specimens, in contrast to the elevated expression reported in hepatocellular carcinoma (HCC) and chronic myeloid leukemia (CML) (23, 46). Moreover, the novelty of our study relies on the observation and definition of the tumor suppressive activity of MITF in PCa. In this context, MITFA upregulation was associated with a reduction in cell proliferation and DNA replication. As occurs in melanoma, the modulation of MITF expression in PCa cells induces the expression of the cell cycle inhibitor p21 but no changes in the cell cycle inhibitor p16 were observed (data not shown). Thus, our results in PCa are in line with the canonical function of MITF in cell cycle progression and proliferation in melanoma (39, 44, 45).

It’s important to highlight that the tissue specific differences in MITF expression among different cancer types suggest that in order to fully comprehend MITF’s role in cancer, its expression and function has to be analyzed in context of each particular cell and tissue type.

CRYAB is a member of the small heat shock protein family that functions as stress-induced molecular chaperone. It inhibits the aggregation of denatured proteins, promotes cell survival and inhibits apoptosis in the context of cancer (47). Paradoxically, CRYAB is highly expressed in some cancer types but decreased in others and in both scenarios an association with cancer progression and prognosis has been reported (25, 26, 28–32, 48–52). In spite of the amount of information regarding the changes in CRYAB expression in cancer, the transcriptional regulation of this chaperone has been poorly explored (48). In this study, we described a novel direct transcriptional regulation of CRYAB by MITF. Although there is no direct nor mechanistic evidence of the MITF-CRYAB transcriptional axis in other cancer types, in melanoma both MITF and CRYAB expression are upregulated by BRAF/MEK-inhibitor treatments (49, 52), suggesting that this regulation can go beyond both prostate cancer scenario. Indeed, we observed that the correlation between MITF and CRYAB is also present in colorectal cancer, but not in breast nor lung cancer (data not shown).

In our study, the MITF-CRYAB transcriptional axis is reduced and exerts tumor suppressive activity in PCa. This is in agreement with the reduced expression of CRYAB observed in PCa patients and its previous consideration as a protective gene against PCa (32). Yet, the exact molecular mechanism underlying the tumor suppressive activity of CRYAB remains to be elucidated.

Importantly, the extensive interrogation of PCa transcriptomes and associated clinical data has led us to propose the transcriptional axis MITF-CRYAB as a potential prognostic biomarker in PCa. The individual expression of CRYAB and MITF has been previously associated with poor prognosis in various tumor types (26, 29–31, 50, 51) and to therapy response in melanoma (53–55). However, our data showing enhanced prognostic potential of the combined signature provides a new and exciting perspective of the functional interaction of these genes in PCa.

Our study endorses the potential of transcriptional deregulation analysis, as either a cause or a consequence of cancer, and its impact to support the discovery of novel cancer related genes and longterm development of novel cancer treatment strategies.

## ACKNOWLEDGMENTS

Apologies to those whose related publications were not cited due to space limitations. The work of V.Torrano is founded by Fundación Vasca de Innovación e Investigación Sanitarias, BIOEF (BIO15/CA/052), the AECC J.P. Bizkaia and the Basque Department of Health (2016111109). The work of A. Carracedo is supported by the Basque Department of Industry, Tourism and Trade (Etortek) and the department of education (IKERTALDEIT1106-16, also participated by A. Gomez-Muñoz), the BBVA foundation, the MINECO (SAF2016-79381-R (FEDER/EU); Severo Ochoa Excellence Accreditation SEV-2016-0644) and the European Research Council (Starting Grant 336343, PoC 754627). CIBERONC was co-funded with FEDER funds. The work of M. Graupera is supported by the MINECO (SAF2014-59950-P). The work of A. Aransay is supported by the Basque Department of Industry, Tourism and Trade (Etortek and Elkartek Programs), the Innovation Technology Department of Bizkaia County, CIBERehd Network and Spanish MINECO the Severo Ochoa Excellence Accreditation (SEV-2016-0644). R. Barrio acknowledges Spanish MINECO (BFU2014-52282-P, Consolider BFU2014-57703-REDC), the Departments of Education and Industry of the Basque Government (PI2012/42) and the Bizkaia County. J. Starková acknowledges the Ministry of Health of Czech Republic AZV NV15-28848A. We are thankful to the Basque Biobank for Research (BIOEF) for the custody and management of human prostate specimens used in this study.

## MATERIALS AND METHODS

### Cell culture and Reagents

Human prostate carcinoma cell lines (PC3) were purchased from Leibniz-Institut DSMZ – Deutsche Sammlung von Mikroorganismen und Zellkulturen GmbH, who provided authentication certificate. The cell line used in this study was not found in the database of commonly misidentified cell lines maintained by ICLAC and NCBI Biosample. Cells were transduced with a modified TRIPZ (Dharmacon) doxycycline inducible lentiviral construct in which the RFP and miR30 region was substituted by *HA-Flag-MITF.* Lentiviral shRNA constructs targeting *PGC1A* (#1-TRCN0000001165 and #2-TRCN0000001166) and *CRYAB* (#1-TRCN0000010822 and #2-TRCN0000010823) were purchased in Sigma and a scramble shRNA (hairpin sequence: CCGGCAACAAGATGAAGAGCACCAACTCGAGTTGGTGCTCTTCATCTTGTTG) was used as control. For *PGC1A* and *CRYAB* shRNAs, Puromycin resistance cassette was replaced by Hygromycin cassette from pLKO.1 Hygro (Addgene Ref. 24150) using *Bam*HI and *Kpn*I sites. Cell lines were routinely monitored for mycoplasma contamination and quarantined while treated if positive. Doxycycline hyclate (Dox) and Puromycin were purchased from Sigma, and Hygromycin from Invitrogen.

### Xenotransplant assays

All mouse experiments were carried out following the ethical guidelines established by the Biosafety and Welfare Committee at CIC bioGUNE. The procedures employed were carried out following the recommendations from AAALAC. Xenograft experiments were performed as previously described (14), injecting 10^6^ cells per condition in two flanks per mouse (Nu/Nu immunodeficient males; 6-12 weeks of age). PC3 TRIPZ-HA-MITFA cells alone or under *CRYAB* silencing were injected in each flank of nude mice and 24 h post-injections mice were fed with chow or doxycycline diet (Research diets, D12100402).

### Patient samples

All samples were obtained from the Basque Biobank for research (BIOEF, Basurto University hospital) upon informed consent and with evaluation and approval from the corresponding ethics committee (CEIC code OHEUN11-12 and OHEUN14-14).

### Molecular assays

Western blot was performed as previously described (14). Antibodies used: HA-Tag (Cell Signalling #3724; dilution 1:10000); MITF (Thermo Fisher Scientific MA5-14146; dilution 1:1000); β-Actin (Cell Signalling #3700; dilution 1:2000); GAPDH (clone 14C10; Cell Signalling #2218; dilution 1:1000); CRYAB (Cell Signalling #45844s; dilution 1:1000).

RNA was extracted using NucleoSpin^®^ RNA isolation kit from Macherey-Nagel (ref: 740955.240C). For patients and animal tissues a Trizol-based implementation of the NucleoSpin^®^ RNA isolation kit protocol was used as reported (14). 1μg of total RNA was used for cDNA synthesis using Maxima™ H Minus cDNA Synthesis Master Mix (Invitrogen M1682). Quantitative Real Time PCR (qRTPCR) was performed as previously described (14). Universal Probe Library (Roche) primers and probes employed are detailed in Supplementary Table 5. *GAPDH* (Hs02758991_g1) housekeeping assay from Applied Biosystems was used for data normalization.

For transcriptomic analysis in PC3 TRIPZ-HA-Flag-MITFA cells, Illumina whole genome -HumanHT-12_V4.0 (DirHyb, nt) method was used as reported (14). A hypergeometric test was used to detect enriched dataset categories.

### Cellular assays

Cell number quantification with crystal violet was performed as referenced (14).

For starvation experiments 100,000 cells per well were seeded in a 6-well plate. Cells were initially plated in 10% FBS media for 24 hours and then the media was changed to FBS free media and left overnight.

Soft agar assays were performed as previously described (14), seeding 5,000 cells per well in 6-well plates.

For BrDu incorporation, cells were seeded on glass cover slips in 12-well plates and after 4 days, cells were incubated with 3 μg mL^-1^ BrDu (Sigma B5002). Cells were fixed with 4% para-formaldehyde, permeabilized with 1% Triton X-100 and incubated with a monoclonal anti-BrDu (MoBU-1) antibody (Invitrogen B35128) at a 1:100 dilution. Images were obtained with an AxioImager D1 microscope (Zeiss). At least three different areas per cover slip were quantified.

### Chromatin Immunoprecipitation

Chromatin Immunoprecipitation (ChIP) was performed using the SimpleChIP^®^ Enzymatic Chromatin IP Kit (Cat: 9003, Cell Signalling Technology, Inc). Four million PC3 cells were grown in 150 mm dishes either with or without 0.5 μg mL^-1^ doxycycline during 3 days. Cells from three 150 mm dishes were cross-linked with 35% formaldehyde for 10 min at room temperature. Glycine was added to dishes during 5 min at room temperature. Cells were then washed twice with ice-cold PBS, and scraped into PBS+PMSF. Pelleted cells were lysed and nuclei were harvested following manufacturer’s instructions. Nuclear lysates were digested with micrococcal nuclease for 20 min at 37°C and then sonicated in 500μl aliquots on ice for 6 pulses of 20 s using a Branson sonicator. Cells were held on ice for at least 1 min between sonications. Lysates were clarified at 11,000 × g for 10 min at 4°C, and chromatin was stored at −80°C. HA-Tag polyclonal antibody (Cat: C29F4, Cell Signalling Technology) and IgG antibody (Cat: 2729, Cell Signalling Technology, Inc), were incubated overnight (4<u>°</u>C) with rotation and protein G magnetic beads were incubated 2hrs (4<u>°</u>C). Washes and elution of chromatin were performed following manufacturer’s instructions. DNA quantification was carried out using a Viia7 Real-Time PCR System (Applied Biosystems) with SybrGreen reagents and primers that amplify the predicted MITFA binding region to *CRYAB* (region 1; For: ttgtttcctcgtagggcttg, Rev: tttcagagccaggagagagc- region 2; For: tctggaatggtgatgtcagg, Rev: attgggtgtggacagaaagc) and *ANGPTL4* (For: gttgacccggctcacaat, Rev: ggaacagctcctggcaatc) as a negative binding control.

### Whole genome gene expression characterization

Whole genome expression characterization was conducted using Human HT12 v4 BeadChips (Illumina Inc.). In brief, cRNA synthesis was obtained with TargetAmp™ Nano-g™ Biotin-aRNA Labeling Kit for the Illumina^®^ System, Epicentre (Cat.Num. TAN07924) and subsequent amplification, labeling and hybridization were performed according to Whole-Genome Gene Expression Direct Hybridization Illumina Inc.’s protocol. Raw data were extracted with GenomeStudio analysis software (Illumina Inc.) in the form of GenomeStudio’s Final Report (sample probe profile).

### Bioinformatic analysis and statistics

#### Database normalization

All the datasets used for the data mining analysis were downloaded from GEO and TCGA. GEO-downloaded data was subjected to background correction, log2 transformation and quartile normalization. In the case of using a pre-processed dataset, this normalization was reviewed and corrected if required. TCGA data were downloaded as upper quartile normalized RSEM count, which was been log2 transformed.

#### Quartile analysis in Disease Free Survival

Patients biopsies from primary tumours were organized into four quartiles according to the expression of the gene of interest in three datasets. The recurrence of the disease was selected as the event of interest. Kaplan-Meier estimator was used to perform the test as it takes into account *right-censoring,* which occurs if a patient withdraws from a study. On the plot, small vertical tick-marks indicate losses, where a patient’s survival time has been right-censored. With this estimator we obtained a survival curve, a graphical representation of the occurrence of the event in the different groups, and a p-value that estimates the statistical power of the differences observed.

#### Correlation analysis

Spearman correlation test was applied to analyse the relationship between paired genes. From this analysis, Spearman coefficient (R) indicates the existing linear correlation (dependence) between two variables *X* and Y, giving a value between +1 and −1 (both included), where 1 is total positive correlation, 0 is no correlation, and −1 is total negative correlation. The p-value indicates the significance of this R coefficient.

#### Statistical analysis

No statistical method was used to predetermine sample size. The experiments were not randomized. The investigators were not blinded to allocation during experiments and outcome assessment. Unless otherwise stated, data analysed by parametric tests is represented by the mean ± s.e.m. of pooled experiments and median ± interquartile range for experiments analysed by non-parametric tests. n values represent the number of independent experiments performed, the number of individual mice or patient specimens. For each independent *in vitro* experiment, at least two technical replicates were used and a minimum number of three experiments were done to ensure adequate statistical power. For data mining analysis, ANOVA test was used for multi-component comparisons and Student T test for two component comparisons. In the *in vitro* experiments, normal distribution was confirmed or assumed (for n<5) and Student T test was applied for two component comparisons. In the statistical analyses involving fold changes, one sample t-test with a hypothetical value of 1 was performed. The confidence level used for all the statistical analyses was of 95% (alpha value = 0.05). Two-tail statistical analysis was applied for experimental design without predicted result, and one-tail for validation or hypothesis-driven experiments.

#### Gene expression array data analysis

first, raw expression data were background-corrected, log2- transformed and quantile-normalized using the lumi R package7, available through the Bioconductor repository. Probes with a “detection p-value” lower than 0.01 in at least one sample were considered expressed. For the detection of differentially expressed genes, a linear model was fitted to the probe data and empirical Bayes moderated t-statistics were calculated using the limma package from Bioconductor. Only genes with differential fold-change (FC) >1.5 or <1.5 and an adjusted p-value < 0.05 were considered as differentially expressed.

### Accession numbers and datasets

#### Primary accessions

The transcriptomic data generated in this publication have been deposited in NCBI’s Gene Expression Omnibus and are accessible through GEO Series accession number GSE114345 (https://www.ncbi.nlm.nih.gov/geo/query/acc.cgi?acc=GSE114345).

#### Referenced accessions

TCGA https://cancergenome.nih.gov/, Grasso *et al.*, GEO: GSE35988 (9); Lapointe *et al.*, GEO: GSE3933 (10); Taylor *et al.*, GEO: GSE21032 (11); Tomlins *et al.*, GEO: GSE6099 (12); Varambally *et al.*, GEO: GSE3325 (13); and Glinsky et al (8).

## AUTHOR CONTRIBUTIONS

LV-J and AM performed the majority of *in vitro* and *in vivo* experiments, unless specified otherwise. NM-M contributed to the in vivo experiments, experimental design and discussion. ARC carried out the bioinformatics and biostatistical analysis. AS-C, MP-V, IH and IA contributed to *in vitro* analysis and provided technical support. IL-V, AS-M, ALI and MU-U provided BPH and PCa samples for gene expression analysis from Basurto University Hospital. MGr carried out microvessel staining and quantifications. JS contributed as supervisor of IH. JDS and RB performed or coordinated (RB) the cloning of *MITFA* in lentiviral vectors. AMMA contributed to experimental design and discussion. AC contributed to experimental design, data analysis and discussion. VT supervised the project, contributed to the experimental design, data generation, analysis and discussion and wrote the manuscript.

